# Specific miRNA-GPCR networks regulate Sox9a/Sox9b activities to promote gonadal renewal in zebrafish

**DOI:** 10.1101/297820

**Authors:** Xinlu Du, Huiping Guo, Ying Zhang, Jiacheng Wu, Minyou Li, Xianxian Hua, Jizhou Yan

## Abstract

Fertility and endocrine function rely on a tightly regulated synchronicity within the hypothalamic-pituitary gonadal (HPG) axis. FSH/cAMP/MAPK/ Sox9 axis signaling and its regulated specific miRNAs are thought to regulate vertebrate gonadal development and sex differentiation, and yet the regulatory networks are largely unknown. Here we construct small RNA and mRNA libraries from sexually matured ovary and testis of zebrafish to identify specific miRNA-target pairs. Integration of Targetscan prediction and *in vivo* induced gene expression highlight four specific miRNAs that conditionally target three G protein–coupled receptor (GPCR) x-Sox9 signaling genes, and implicate two regulatory circuits of miR430a-Sox9a in the testis and miR218a-Sox9b in the ovary. Co-injected Sox9a-miR430a mixture increases the proportion of spermatogonia but degenerates primary oocyte, while Sox9b-miR218a mixture induces renewal of ovarian follicles. Co-immunoprecipitation and mass-spectrometry analyses further reveal that miR430a and Sox9a synergistically activate testicular PKC/Rock1 signals while miR218a and Sox9b constrict ovary PKC/PI3K/Rock1 signaling. These results clarify specific miRNAs-GPCR regulatory networks of Sox9a/Sox9b switch, and also provide mechanistic insight into gonadal rejuvenation and plasticity.

## Introduction

Although cell phenotypic changes often occur in stem cell-based differentiation, dedifferentiation or transdifferentiation, both stem cells and differentiated cells are able to reversibly dedifferentiate and transform into cells of different lineages under certain conditions. This phenomenon, called environmentally guided cell plasticity is the basis of tissue homeostasis, tissue regeneration, cancer cell recover (Galliot & Ghila, 2010), and sex reversal (Sun, Zhang et al., 2013). One excellent system to study cell plasticity is the vertebrate gonad, where distinct populations of somatic and germ stem cells give rise to different sex-dependent morphotypes: male testiculogenesis and female folliculogenesis (Chen, Zheng et al., 2013, Gassei & Schlatt, 2007). The early gonad is an undifferentiated primordium composed of bipotential somatic stem cells: precursors for supporting cells and steroid-secreting cells. When the primordial germ cells (PGCs) reside in the gonadal ridge, the supporting cell precursors develop into either testis-specific Sertoli cells or ovary-specific follicle (granulosa) cells. Meanwhile the immigrating mesonephric cells give rise to peritubular myoid (PM) cells, endothelial cells that form the male-specific vasculature, and fetal steroidogenic-Leydig cells (or ovarian theca cells in female) (McClelland, Bowles et al., 2012, Wilhelm, Palmer et al., 2007). In parallel to the differentiation of somatic precursors, PGCs differentiate into oogonia in ovarian follicles or into spermatogonia in Sertoli cells-dominant testis cord.

Histologically, testiculogenesis with cord formation and Leyding cell differentiation supports spermatogenesis while folliculogenesis accompanies ootidogenesis, where the ovarian follicle surrounding the oocyte develops from a primordial follicle to a preovulatory one. There is a close anatomical relationship between the development of the genital ridge and the hormone excretory system during early ontogeny of all vertebrates, including fish. In both teleosts and mammals the testis contains Sertoli and Leydig cells in addition to germ cells, and the ovary consists of the thecal cells and granulosa cells surrounding the ovum. The gonadal sex differentiation involves the same sequential cellular events for spermatogenesis and folliculogenesis, in which ovarian development is the default pathway. Without Sox9-related signaling intervention, the bipotential gonads follow the ovarian pathway (McClelland et al., 2012, Sun et al., 2013).

An important difference is that the mammalian gonads are terminally developed into either testis or ovary by Sry-Sox9 genetic intervention, while fish gonads often retain the ability to change, making them sequential hermaphrodites. Despite the vast diversity of primary sex-determining mechanisms in vertebrate species, Sox9-related signals appear to be a conserved mechanism to switch on testis-differentiation pathway, where the expressions of *sox9* and its downstream genes (*amh* and *cyp19a1*) are proposed to be regulated by gonadotropins and its second messenger cAMP through MAPK signaling pathways (Kobayashi, Chang et al., 2005, Taieb, Grynberg et al., 2011). Intricately, activation of Sox9-MAPK pathways are also linked to chondrocyte differentiation (Murakami, Kan et al., 2000). Accumulating evidence implicates versatile functions of Sox9 in cell fate specification, stem cell biology, human diseases and tissue regeneration (Jo, Denduluri et al., 2014). Therefore, identifying partner factors, signaling pathways, post-transcriptional modifications, and integrative signaling pathways could facilitate us to better understand Sox9’s versatility in cell fate switch and adaptive plasticity.

Zebrafish has two co-orthologs of *sox9* on its duplicated chromosomes: *sox9a* and *sox9b*. The pair of *sox9* genes show distinct and overlapping functions in zebrafish embryonic development (Rodriguez-Mari, Yan et al., 2005, Yan, Willoughby et al., 2005). In the adults, *sox9a* is expressed in many tissues including testis, whereas *sox9b* expression is restricted to previtellogenic oocytes of the ovary (Chiang, Pai et al., 2001). Correspondingly two Sox9 downstream genes, *amh* and *cyp19a1a*, are also respectively expressed in the juvenile testis and ovary (Rodriguez-Mari et al., 2005). Therefore, zebrafish bipotential gonad is a unique system to explore more informative *sox9* axis signaling pathways controlling gonadal sexual fate. MicroRNAs (miRNAs) represent a second regulatory network that regulates gene expression in various biological processes by inducing degradation or translational inhibition of their target mRNAs (Tao, Sun et al., 2016). Based on miRNA expression in the gonads at the time of sexual differentiation, a number of miRNAs and the potential targets are identified. Some specific miRNAs show regulatory activities in gonad development and/or sex determination (Kang, Cui et al., 2013, Xiao, Zhong et al., 2014). Even though miR-124 was reported to be sufficient to induce the repression of both SOX9 translation and transcription in ovarian cells (Real, Sekido et al., 2013), no the specific miRNA-*sox9* axis regulatory networks have been identified so far (Tao et al., 2016). In this study we performed genome-wide search for specific miRNA-target pairs, and investigated physiological interaction effects between specific miRNAs and the target genes. Particularly, we focused on specific miRNA-Sox9a/Sox9b networks in regulation of testiculogenesis and folliculogenesis.

## Results

### Identification of genes related to sex-specific fate during the process of gonadal differentiation maturation

To identify genes functioning in folliculogenesis and testiculogenesis, we dissected the freshly matured gonads at 5mpf (months post-fertilization), and separated the ovary and testis for RNA deep sequencing analyses. Out of 26,459 genes (Table S1), 25,609 genes were expressed in either or both sex gonads. 18,993 genes exhibited 2 fold or more expression changes (Table S2), and 729 genes were significantly differentially expressed in testis or ovary (P˂0.05) (Table S3), including 421 ovary-, and 308 testis-highly expressed (Table S4). As the predominant expression of these induced genes synchronized with the reproductive growth cycle (testicular or follicle differentiation), we categorized 729 biased genes as sex-preferential effectors and/or regulators.

Our initial goal was to clarify the proposed gonadotropins-cAMP-MAPK-Sox9 signaling pathways during sex gonadal differentiation, Gonadotropins including follicle-stimulating hormone (FSH) and luteinizing hormone (LH), are glycoprotein peptide hormones. Gonadotropin receptors are coupled to the G-protein system. We searched the two transcriptome datasets for GPCR*-sox9* axis genes. These included 92 GPCR, 35 cAMP, 34 MAPKs, and 21 p53 orthologs as well as 30 sex determinant genes. The vast majority of the GPCR*-sox9* axis genes (201/212) did not show significant differential expression between testis and ovary (Table S5). This result conformed to the phenomena that miRNAs only modestly downregulate the mRNA level of their target genes (Selbach, Schwanhausser et al., 2008).

### Identification of specific miRNAs and GPCR*-sox9* axis target genes for gonadal differentiation

To identify specific miRNAs-target pairs in reproductive cycle, we analyzed two small RNA libraries, which were simultaneously constructed from the same ovary or testis total RNA samples as the mRNA libraries were made. After aligning the small RNA sequences with the miRBase (zebrafish miRNA database), bioinformatic analysis identified 350 unique mature miRNAs, and 346 miRNA precursors. 314 miRNAs were found in either of the two libraries, of which 51 miRNAs showed 2-fold higher transcription in testis (refer to testis miRNAs) and 106 miRNAs were preferentially transcribed in ovary (ovary miRNAs).

Using TargetScan to search for the target pairs between 157 sex-biased miRNAs and 209 GPCR*-sox9* axis genes, we found that 97 miRNAs targeted 183 GPCR-Sox9 axis genes, including those multiple and cross targets (Table S7). For example *sox9a* was targeted by 25 miRNAs and *sox9b* was targeted by 20 miRNAs.

### Four specific miRNAs represent putative regulators in *GPCR-sox9*-axis

To derive a short-list of specific miRNAs as putative regulators in GPCR-*sox9* axis, we selected the representative miRNAs-target pairs according to the criteria in Table 1. Combined with the expression pattern of endogenous pre-miRNAs (Fig.S1), four potentially interesting miRNAs were accordingly selected as the most overrepresented miRNAs: miR430a (multiple splicing isoforms and 17 multiple targets in GPCR*-sox9* axis of signaling cascades, not directly targeting *sox9*), miR218a (50 targets, not directly targeting *sox9* but targeting *vasa*), miR734 (95 targets, directly targeting *sox9a*), and miR141 (72 targets, directly targeting *sox9b*) (Table S7).

**Table 1.**
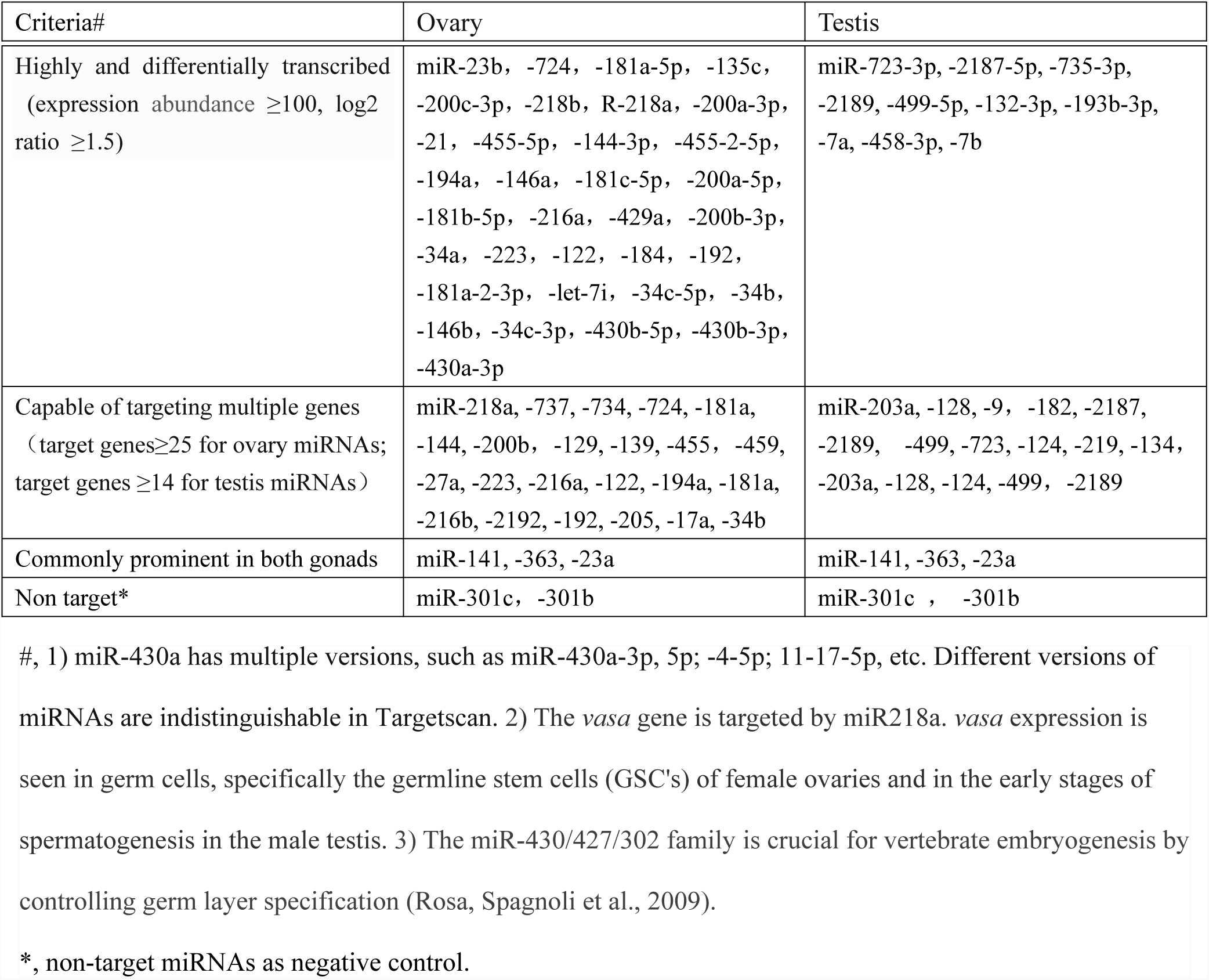
A short list of overrepresented miRNAs

| Criteria# | Ovary | Testis |
| --- | --- | --- |
| Highly and differentially transcribed<br>(expression abundance $\geq 100$, log2<br>ratio $\geq 1.5$) | miR-23b, -724, -181a-5p, -135c,<br>-200c-3p, -218b, R-218a, -200a-3p,<br>-21, -455-5p, -144-3p, -455-2-5p,<br>-194a, -146a, -181c-5p, -200a-5p,<br>-181b-5p, -216a, -429a, -200b-3p,<br>-34a, -223, -122, -184, -192,<br>-181a-2-3p, -let-7i, -34c-5p, -34b,<br>-146b, -34c-3p, -430b-5p, -430b-3p,<br>-430a-3p | miR-723-3p, -2187-5p, -735-3p,<br>-2189, -499-5p, -132-3p, -193b-3p,<br>-7a, -458-3p, -7b |
| Capable of targeting multiple genes<br>(target genes $\geq 25$ for ovary miRNAs;<br>target genes $\geq 14$ for testis miRNAs) | miR-218a, -737, -734, -724, -181a,<br>-144, -200b, -129, -139, -455, -459,<br>-27a, -223, -216a, -122, -194a, -181a,<br>-216b, -2192, -192, -205, -17a, -34b | miR-203a, -128, -9, -182, -2187,<br>-2189, -499, -723, -124, -219, -134,<br>-203a, -128, -124, -499, -2189 |
| Commonly prominent in both gonads | miR-141, -363, -23a | miR-141, -363, -23a |
| Non target* | miR-301c, -301b | miR-301c, -301b |

#, 1) miR-430a has multiple versions, such as miR-430a-3p, 5p; -4-5p; 11-17-5p, etc. Different versions of miRNAs are indistinguishable in Targetscan. 2) The *vasa* gene is targeted by miR218a. *vasa* expression is seen in germ cells, specifically the germline stem cells (GSC's) of female ovaries and in the early stages of spermatogenesis in the male testis. 3) The miR-430/427/302 family is crucial for vertebrate embryogenesis by controlling germ layer specification (Rosa, Spagnoli et al., 2009).
\*, non-target miRNAs as negative control.

### Interaction modes between specific miRNAs and GPCR*-sox9* targets in testis and ovary

To investigate the physiological interactions between four specific miRNAs and GPCR*-sox9* axis targets, we devised an *in vitro* gonadal microinjection strategy to enforce exogenous expression of Sox9a/Sox9b and four specific miRNAs in the gonads, and evaluated their effects on the expression of GPCR*-sox9* axis targets (Table S8). When the Targetscan prediction and RT-PCR examinations were integrated together, four interaction modes emerged (Table 2): 1) a negative specific miRNA-target pair: a predicted target was downregulated in both gonads by a specific miRNA; 2) a positive specific miRNA-target pair: a predicted target was upregulated in both gonads by a specific miRNA; 3) a conditionally specific miRNA-target pair: a predicted target was upregulated by a specific miRNA in one sex gonad but downregulated in the other gonad; and 4) an indirect specific miRNA-target: a gene without specific miRNA binding sites was downregulated or upregulated. Totally 92.3% (12 of 13) of predicted targets of GPCR*-sox9* genes were correctly scored as direct targets in testis and/or ovary by at least one of four specific miRNAs. Fig.1 showed crossing alignment between four specific miRNAs and 15 putative target genes.

**Table 2.**
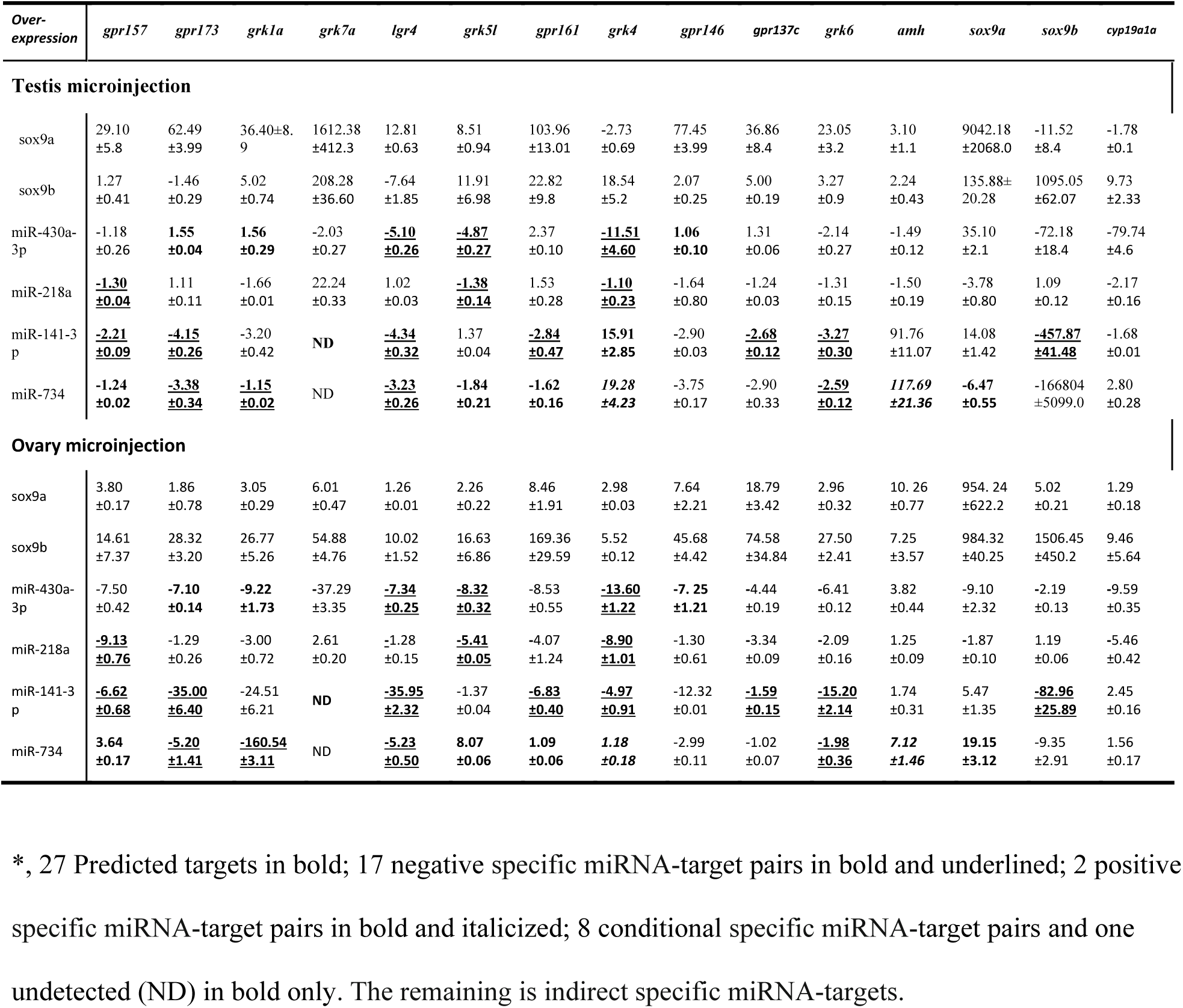
Interactions between specific miRNAs and GPCR-Sox9 axis^*^

**Fig 1.**
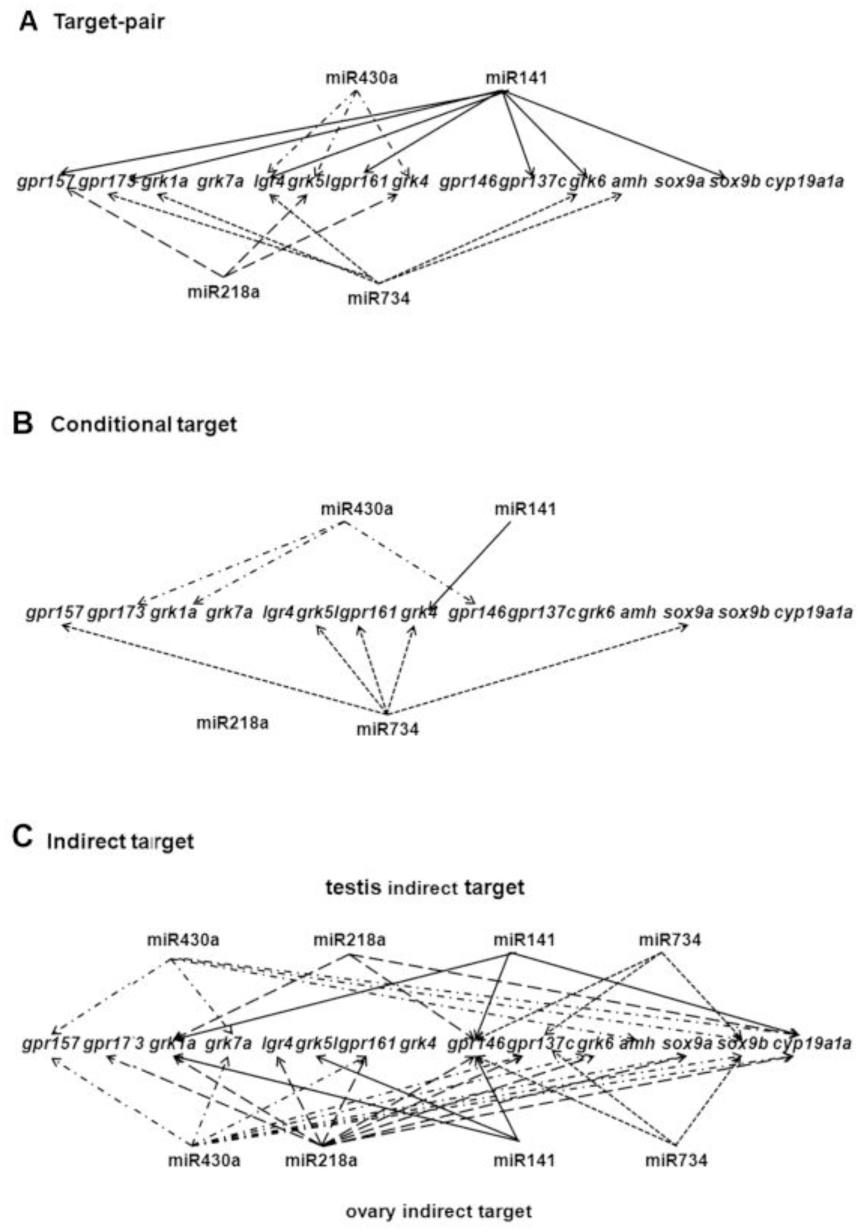
Interaction modes between four specific miRNAs and GPCR-Sox9a/b targets.

### Specific miRNAs-GPCR network regulate Sox9a/Sox9b signaling at post-transcription and protein level

Previous study has identified several *sox9* downstream genes (*cyp19a, cyp19b, amh)* as potential targets of miR430 family during sexual transformation of the rice field eel (Gao, Guo et al., 2014). Here we found that miR734 and miR141 formed target-pairs or conditional target pairs with *sox9a, amh* and/or *sox9b*, while miR430a and miR218a only showed indirect interactions with these genes and *cyp19a1a* (Fig.1). Referred to *sox9a* and *sox9b* transcription, increased miR141 and miR734 showed similar regulatory functions as miR430a (Table S8). Additionally three miR430a direct target genes (*lgr4*, *grk5l, grk4*) were also pair-targets or conditional pair-targets for miR141, miR734 and miR218a (Fig.1). Thus, the four miRNAs may coordinate to regulate Sox9a/Sox9b activity through GPCR-signaling networks.

We then investigated the regulatory modes between the four specific miRNAs and two Sox9 isoforms. Sustained Sox9b expression increased the transcription of four pre-miRNAs in both ovary and testis, whereas enforced Sox9a only induced pre-miR430a transcription in testis (Fig.S2). Similarly, overexpressed Sox9a and Sox9b activated all tested GPCR-*sox9* genes in the ovary whereas a few genes were suppressed in testis.

When mature miR430a oligos were injected into the testis, *sox9a* transcript was increased but *sox9b* transcript was strongly reduced. In the ovary, elevated miR430a somewhat reduced transcripts of *sox9a, sox9b* and *cyp19a1a*. In contrast to miR430a-Sox9a reciprocal activation in the testis, miR218a downregulated transcripts of *sox9a* and *cyp19a1a* but modestly upregulated *sox9b* transcript in both testis and ovary. These results implicated two transcriptional regulatory circuits: miR430a-Sox9a, and miR218a-Sox9b. Relative to miR218a-Sox9b’s extensive and ovary-biased regulation, miR430a-Sox9a synergic regulation was specific in the testis. When we constructed a GFP reporter for 3’UTR of *sox9a* and *sox9b*, and carried out GFP reporter assays in zebrafish embryonic fibroblast cells (Pac2), we found that miR430a and miR218a not only regulates the two Sox9 genes’ transcription, but also confines the two proteins’ subcellular distribution (Fig.S4, S5).

### Gonadal microinjection of specific miRNAs-Sox9 mixture cause renewal of gonocytes

To evaluate two miRNA-Sox9 regulatory circuits and its role in the gonadal differentiation, we conducted a combinatory microinjection into mature gonads (5mpf), and then compared cellular events of somatic cells and germline cells (Fig.2A, 2B). Enforced Sox9a-miR430a mixture expression increased the proportion of mitotic condensed chromatin nuclear cell (Cn) and undifferentiated gonocytes (Gc), and increased degeneration of primary oocytes (Fig.2Aa, d). Co-injected Sox9a-miR141-miR734 cocktails increased proportion of spermatocytes (Sc), and also induced ovary-testis transition (indicated by oocyte degeneration and multinucleate cell generation, Fig.2Ab, 2Ae). Conversely, enforced Sox9b-miR218a expression increased each stage of follicles (oocytogenesis), but reduced testis Cn and Sc with increase of deforming Gc (Fig. 2Ac, 2Af). We didn’t found consistent roles of other Sox9a/ Sox9b-miRNAs mixtures in testiculogenesis and folliclulogensis (Fig.2C, Fig.S3).

**Fig 2.**
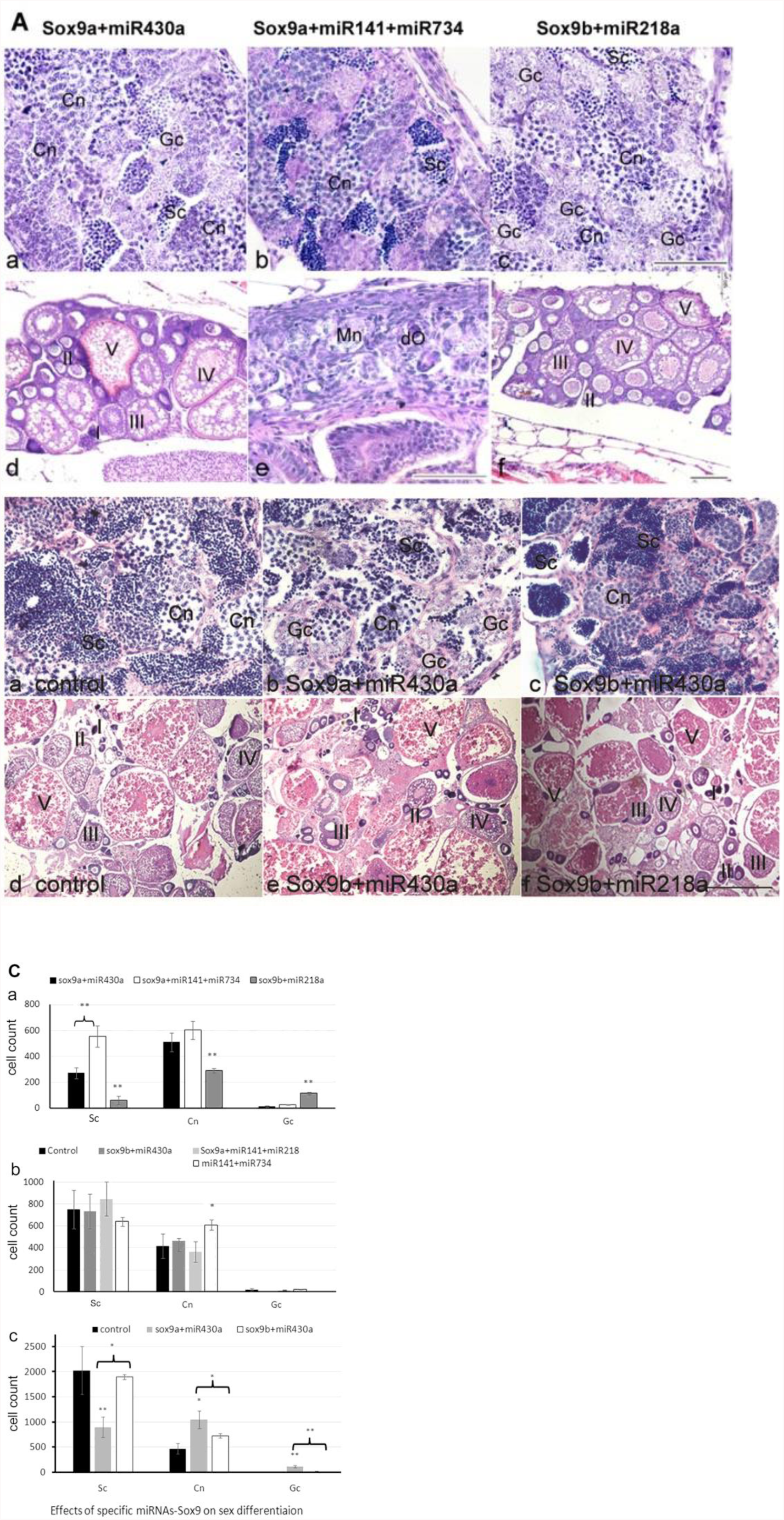
Effects of exogenous Sox9a/Sox9b coupled specific miRNAs on testiculogenesis and folliculogenesis. (A) Histological observations on testiculogenesis and folliculogenesis in mature zebrafish (5 mpf). Based on cell morphology, the germline cells were classified into three categories: gonocytes (Gc), mitotic condensed chromatin nuclear cell (Cn), and mature sex gamete (Gm). Gc includes spermatogonia, oogonia or other undifferentiated PGCs. (Cn) includes spermatocytes, or primary oocytes (pOc), Gamete includes spermatids or spermocyte (Sc), secondary oocyte within five stages of follicles (I-V). Degenerating oocytes (dO) were indicated by arrow. Some enlarged gonocytes (indicated by arrow head) looked like early perinucleolar oocytes (or regarded as deforming gonocytes). (B) Histological observations on testiculogenesis and folliculogenesis in old fish (1.5 years old). a-c show testiculogenesis; d-f show various stages of follicluogenesis: stage Ia, Ib, II, III, and IV oocytes. (C) Statistical analyses on cell count of germ cells at various stages of spermatogenesis and oogenesis. The proportion of each category was compared among different specific miRNA-Sox9a/b treatments for 5mf fish (a, b) and 1-1.5 years old fish (c). * (p<0.05), ** (p<0.01). Scale bar =50 μm (testis) and 500 μm (ovary).

Since Sox9a, Sox9b and pre-miRNAs transcripts were differentially transcribed in the testis and ovary (Fig.S1), and naturally reduced with age, we posited that exogenous specific miRNAs-coupled Sox9a or Sox9b may induce certain “rejuvenation" effects on the old gonads. To explore this possibility, we tested 1.5-2 years old of zebrafish. As expected, Sox9a-miR430a mixture significantly increased the proportion of Gc and Cn whereas Sox9b-miR218a mixture increased all stages of follicles. Sox9b+miR430a increased the testis Cn and Sc (Fig.2Bc, 2Cc) as well as the primordial follicles (Fig.2Be). Compared to the injection into the young fish (5 months olds) (Fig.2A), Sox9b-miR218a mixture (Fig.2A–2B) more significantly increased the ratio of the primordial follicles in the old fish.

To further test the rejuvenation effects, we performed the same specific miRNA-Sox9a/b microinjection into the old gonads of *vasa* promoter-driven GFP transgenic medaka. Germ cell-specific gene, *vasa*, is most abundantly expressed in mitotic germ cells (oogonia and spermatogonia) and reduced in meiotic germ cells (Tanaka, Kinoshita et al., 2001, Yuan, Li et al., 2014). As shown in Fig.S3B, miR430a-Sox9a increased the testis GFP-positive cells; miR430a-Sox9b increased the ovarian primordial follicles; and miR218-Sox9b increased all stages of follicles (mostly of primordial follicles). These results reaffirmed that miR430a-Sox9a coordination is sufficient to induce spermatogenesis while miR218a-Sox9b could promote renewal of ovarian follicles.

### MiR430a and miR218a conversely regulate chemical modifications of Sox9a/Sox9b proteins

Since *grk4* and *grk5l* encode the members of G protein-coupled receptor kinase subfamily, our proposed specific miRNA-GPCR networks may regulate Sox9a and Sox9b protein modifications. Presumably covalent modifications in Sox9a and Sox9b proteins would change the molecular-weight sizes and/or band patterns. Immunoblotting analyses showed that specific Sox9a and Sox9b antibodies recognized several bands, which molecular-weight sizes were bigger than the predicted; and the band patterns were changed after cotransfections of miR430a or miR218a (Fig.3A). In the transfected Pac2 cells, the presumed protein modifications (Fig.3.B) were not significant as those occurred in the microinjected gonadal tissues.

**Fig 3.**
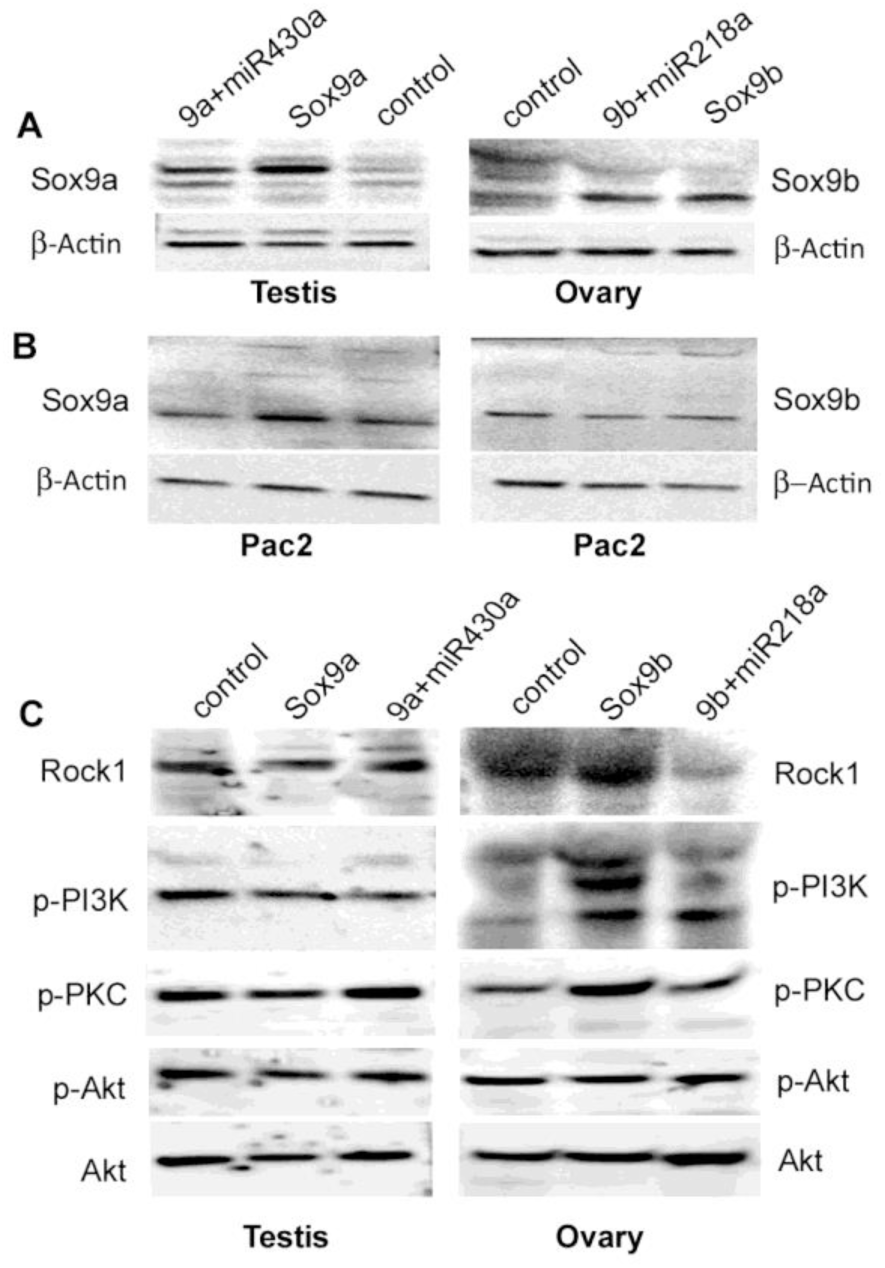
Immunoblotting analyses of endogenous and exogenous Sox9a/Sox9b protein modifications. (A) The testes were microinjected with pSox9a-myc (Sox9a) or pSox9a-myc and miR430a mix (9a+miR430a); the ovaries were injected with pSox9b-myc (Sox9b) or pSox9b-myc and miR218a mix (9b+miR218a). (B). In the *in vitro* cultured Pac2 cells were transfected with pSox9a-myc, pSox9a-myc/miR430a mix, pSox9b-myc, pSox9b-myc/miR218a mix, or empty-control. (C). Sex gonads were microinjected as described in (A). The expression of each signaling component was examined. Relative signal intensity was calculated compared to total Akt.

Co-immunoprecipitation (Co-IP) coupled-mass spectrometer analysis indicated several reverse changes in covalent modifications (Table 3). First, enforced Sox9a-injected gonadal tissues showed enriched Sox9a peptide phosphorylation, acetylation and ubiquitination. On co-injection with miR430a, Sox9a phosphorylations and ubiquitination decreased, but oxidation increased. Meanwhile phosphorylation, ubiquitination and carbamidomethyl at Sox9b and Sox8b peptides were increased by enforced miR430a-Sox9a mixture. Second, enforced Sox9b induced extensive Sox9b acetylation while addition of miR218a retained Sox9b acetylation, but increased oxidation and phosphorylation at Sox9b, Sox9a and Sox8a peptides. Third, both Sox9a and Sox9b peptides were highly acetylated, and mostly accompanied with other modifications. Statistical analysis indicated that the putative enzymes responsible for significantly enriched phosphorylations, and acetylation, included phosphatidylinositol 3-kinase, rho-associated protein kinase, mitogen-activated protein kinase-activated protein kinase 2a, protein kinase C, novel protein similar to vertebrate carnitine acetyltransferase (CRAT), n-alpha-acetyltransferase 35, and acetyl-CoA acetyltransferase. We then verified whether the identified kinases linked to our proposed miR430a/miR218a-GPCR-Sox9 networks. Enforced Sox9b increased Rock1, p-PI3K and p-PKC in the ovary while co-injected miR218a suppressed Rock1 and p-PKC, partially suppressed p-PI3K, and increased p-Akt, suggesting that miR218 could modulate Sox9b activations (Fig.3C). In the testis, Sox9a did not activate Rock1, p-PI3K and p-PKC; However, Sox9a and miR430a mixture activated p-PKC and Rock1signaling.

**Table 3.** CoIP-Mass spectrometer analyses on chemical modifications in Sox9a and Sox9b peptides

| Enforced | Sequence | Matched | Modifications * | Putative enzymes |
| --- | --- | --- | --- | --- |

| expressi<br>on# |  | protein |  |  |
| --- | --- | --- | --- | --- |
| Sox9a | NGQSESEDGSEQTHI<br>SPNAIFK | Sox9a | 3xPhospho [S4; S6;<br>S10]; 1xAcetyl [K22] | Phosphatidylinositol<br>3-kinase, Rho-associated<br>protein kinase,<br>Serine/threonine-protein<br>kinase SBK1 ,<br>Tyrosine-protein kinase<br>BAZ1B (Fragment) |
|  | TLGKLWRLLENEVEK | Sox9a | 1xAcetyl [T/K];<br>1xPhospho [T1]; 1xGG<br>[K14] |  |
| Sox9a+<br>miR430a | NKPHVKRPMNAFM<br>VWAQAARR | Sox9a | Acetyl [N];<br>1xOxidation [M9] | Novel protein similar to<br>vertebrate carnitine<br>acetyltransferase<br>(CRAT) ;<br>N-alpha-acetyltransferas<br>e 35,<br>Mitogen-activated<br>protein kinase-activated<br>protein kinase 2a. |
|  | ADLKR | Sox9a,Sox9b, | 1xAcetyl [N-Term] |  |
|  | TDLPVCSKADLKR | Sox9b | 1xPhospho [T1];<br>1xAcetyl [T/K] |  |
|  | VSGSGKSKPHVK | Sox9b,Sox8b | 1xPhospho [S7];<br>1xAcetyl [N-Term];<br>2xGG [K6; K8] |  |
| Sox9b | VSGSGKSKPHVK | Sox9b | 1xAcetyl [K6] | Acetyl-CoA<br>acetyltransferase |
|  | LRVQHKK | Sox9b | 2xAcetyl[K6; K7] |  |
|  | DAVSQVLK | Sox9b |  |  |
| Sox9b+<br>miR218a | RPMNAFMVWAQAA<br>RRK | Sox9a,<br>Sox9b,Sox10,<br>Sox8a | 2xOxidation [M3; M7] | Acetyl-CoA<br>acetyltransferase,<br>Protein kinase C. |
|  | DAVSQVLK | Sox8a,Sox9b |  |  |
|  | VSGSGKSKPHVKRP<br>MNAFMVWAQAAR | Sox9b, Sox8,<br>Sox9a | Acetyl [K]; 1xPhospho<br>[S7]; 2xOxidation<br>[M15; M19] |  |
|  | ADLKRER | Sox9b | 1xAcetyl [K4] |  |
|  | VNGSSKNKPHVKRP<br>MNAFMVWAQAAR | Sox9a | Acetyl [K];<br>1xOxidation [M19] |  |

#, Sex gonads from 20 Fish (2 months post fertilization, uncertain male or female) were divided four groups and separately injected by 1) Sox9a-Myc plasmid; 2) Sox9a-Myc plasmid+miR430a+miR141; 3)Sox9b-Myc plasmid; and 4) Sox9b-Myc plasmid+miR218a. Two more injections were repeated in the first week. Two weeks later, the gonads were collected for co-immunoprecipitation with Sox9a-Myc antibody mix (group 1

Although specific antibodies are required to verify the exact modifications in Sox9a and Sox9b proteins, together with the results shown in Fig.3, Fig.S2, Table2 and Table 3, we can make following conclusions: 1) Sox9b up-regulated GPCR genes (*lgr4*, *grk5l, grk4*), and extensively activate PKC/PI3K/Rock1 signaling pathways and Sox9a/b modifications while miR218a could restrain such activations possibly by phosphorylation of Sox9b through PI3K-pAkt pathway. 2) In the testis, miR430a and Sox9a reciprocally regulate GPCR genes (*lgr4*, *grk5l, grk4*) and activate p-PKC-MAPK/Rock1 pathways to repress Sox9b activities.

## Discussion

Inducing cell phenotypic conversions usually require multi-purpose signaling inputs and major remodeling of gene transcription (Zhang, Stokes et al., 2011). Specific miRNAs and targets have been identified to account for cell identity changes of skin stem cells (Zhang et al., 2011), mesenchymal progenitors (Huang, Zhao et al., 2010), and cardiomyocytes (Xin, Olson et al., 2013). Functionally phenotypic conversion of sex gonadal cells appear more intriguing, because it involves coordination of multiple somatic and germline lineages in order to successfully commit testiculogenesis or folliculogenesis. As scientists are trying to translate the information gained from basic scientific studies on ovarian organogenesis to the development of *in vitro* strategies to derive stem-cell based ‘prosthetic ovary’ (Truman, Tilly et al., 2017), an attractive possible alternative step in this process is discovery of new ways to drive oogenesis *in vivo*. In the present study, we find that specific miRNAs-GPCR networks could modulate and renew spermatogenesis and folliculogenesis.

First, we provide molecular basis for miRNAs-posttranscriptional regulation of Sox9 switch in the gonadal sex differentiation. In mammals, sex differentiation is controlled by two groups of sex-determining genes that promote one gonadal sex and antagonize the opposite one (Rodriguez-Mari et al., 2005). Zebrafish has no sex chromosome but has two isoforms of *sox9* gene: *sox9a* and *sox9b* (Chiang et al., 2001). Our in vitro cell transfections and in vivo gonadal microinjections indicated that miR430a-Sox9a network could induce spermatogenesis while miR218a-Sox9b network promote folliculogenesis. We established miR430a/miR218a-GPCR-Sox9a/Sox9b switch regulatory model. In this model, miR430a and miR218a convergently target *lgr4, grk5l and grk4* GPCR genes, and conditionally regulate PI3K/ PKC/Rock signaling pathways and Sox9a/Sox9b activities in two sex gonads. MicroRNA-340 inhibits invasion and metastasis of cancer cells by downregulating ROCK1 in (Maskey, Li et al., 2017), however miR430a and Sox9a mixture coordinately activate Rock1 and PKC pathways to induce spermatogenesis. In the ovary Sox9b and miR218a together could restrain PI3K/ PKC/Rock signaling to increase primordial follicle reserve. This repression is important because over-activation of PI3K/ mTORC1 signaling pathways could accelerate depletion of primordial follicles, and lead to premature activation of the entire pool of primordial follicles (Adhikari, Zheng et al., 2010, Reddy, Liu et al., 2008).

GPCRs (G protein-coupled receptors) were found to play crucial roles in maintenance of planarian germ cell plasticity (Saberi, Jamal et al., 2016). We identified three putative GPCR molecules and several modifier enzymes responsible for Sox9a and Sox9b posttranscriptional modifications (Table 3). Given that Lgr4, Grk5l and Grk4 could respectively link to gonadotropins(Hsu, Liang et al., 1998), mTORC1 activity (Burkhalter, Fralish et al., 2013), as well as SUMO modification (Yu, Yin et al., 2017), our finding not only supplements the previously described composition and function of G protein-coupled receptor signalsomes (Bar Oz, Kumar et al., 2016, Luttrell, 2005, Spiegelberg, 2013), but also clarifies the signal transduction of gonadotropins/GPCR/MAPK/ Sox9a/b switch pathways.

Second, our analysis on the global mRNA-miRNA expression profiles that concur in the same sex gonads represents the first effort to systematically identify the gonadotropins-GPCR-Sox9 regulatory networks in testiculogenesis and folliculogenesis. We extend the previous finding that miRNAs could target multiple genes and form a complex miRNA regulatory network (Tao et al., 2016), and summarize four interaction modes between specific miRNAs and the target genes, *i.e*., downreguatory target-pair, upregulatory target-pair, conditionally regulatory target-pair, and indirectly regulatory target-pair.

Previous studies have revealed dual regulatory module of miRNAs from repression to activation at transcriptional and translational levels (Vasudevan, Tong et al., 2007). The miR-430 allows the post-transcriptional activation of *nanos*1 in germline cells, but induces deadenylation and translational repression in somatic cells (Mishima, Giraldez et al., 2006). Now we present the other mechanism that miR430a could modulate expression and intercellular localization of Sox9a and Sox9b proteins through chemical modifications, such as phosphorylation, acetylation, oxidation and ubiquitination (Table 3). It has been found that these types of posttranscriptional modifications could modulate the stability, intracellular localization, and the overall activity of Sox9 (Bar Oz et al., 2016, Jo et al., 2014, Xu, Paige et al., 2010). Finally, we initiated gonadal injections of the specific miRNA-target cocktail to rejuvenate gonadal reserve. Replacement therapy with hormones, and stem cells, blood from aged male, and blood mononuclear cells have been tested to promote ovarian follicle dynamics, spermatogenesis, and tissue rejuvenation (Bukovsky, 2015, Moss, Crosnoe et al., 2013, Niikura, Niikura et al., 2010, Tilly & Telfer, 2009, Zouboulis & Makrantonaki, 2012). However our knowledge of how these factors and cells repair aged reproductive functions still remains limited. Although adult mouse ovaries retain proliferative germ cells resembling male spermatogonial stem cells *in vitro* (Tilly & Telfer, 2009), it is desirable to induce oogenesis under physiological conditions. In the present study, we found that gonadal injection of miR430a-Sox9a mixture could increase testicular gonocytes, and miR218a-Sox9b mixture could increase primordial follicles in old ovaries. Although it is unknown about the source origins from proliferation of reserved stem cell or dedifferentiation of adult gonadal cells, miR430a-Sox9a and miR430a-Sox9b at least could expand the testis spermatogonia reserve and ovarian follicle reserve respectively. If equivalent efficacy can be found in human gonads, miRNA-based testis rejuvenation and prevention of premature ovarian failure may one day be realized. The miRNA-target pair data reported here will help identify more potential therapeutic targets and promote to develop new therapeutic interventions for aging-related ovarian failure and age-related testicular regression.

## Materials and methods

### Fish husbandry

Zebrafish and *vasa*-GFP transgenic medaka were bred and maintained as previously described (Li, Guan et al., 2013, Wang, He et al., 2012, Yuan et al., 2014). All experiments with zebrafish and medaka were approved by the laboratory animal care and use committee of Shanghai Ocean University, and performed in accordance with “Guide for the Care and Use of Laboratory Animals” (NIH).

### Gonadal microinjection

A simple and feasible method for body surface injection into sex gonad has been described (personal data). Briefly, we dissected ten adult zebrafish (five each sex), measured the size ratio of gonad to the body, and figured out the integrated surface projection of the sex gonads, and determined the injection area. Based on the relative size and location of the sex gonads, injection site was about 1 mm upward from the intersect between pectoral fin upper horizontal and the front end vertical line of spelvic fin. By this way overexpression plasmids (100ng), and/or specific miRNAs (20pmol each oligos) (Table S10) were microinjected into sex gonads with lipohigh transfection reagent (Sangon Biotech, Shanghai). Microinjection efficiency was confirmed by western blotting analyses of enforced Sox9a-Myc and Sox9b-Myc (Fig.S6).

### Data Availability Statement

All relevant data are within the paper and its supporting information files. All RNA sequencing data are available at the NCBI short read archive: under study number PRJNA293388 (SRP062685) for mRNA transcriptome; under study number PRJNA294493 (SRP063086) for small RNA.

## Acknowledgements

We thank Ying Peng for her technical assistance in immunoblotting analyses. This work was supported in part by Shanghai Universities First-class Disciplines Project of Fisheries to JY, and National Natural Science Foundation of China (31672700) to ML. The funders had no role in study design, data collection and analysis, decision to publish, or preparation of the manuscript.

## Author contributions

Conceived and designed the experiments: JY. Performed the experiments: XD, HG, YZ, JW. Analyzed the data: JY, XH, XD, HG, YZ, JW. Wrote the paper: JY, XH, ML.

